# Mutations in *rv0678* confer low-level resistance to benzothiazinone DprE1 inhibitors in *M. tuberculosis*

**DOI:** 10.1101/2022.06.29.498178

**Authors:** Nicholas C. Poulton, Zachary A. Azadian, Michael A. DeJesus, Jeremy M. Rock

## Abstract

Tuberculosis (TB) is the leading cause of death from any bacterial infection, causing 1.5 million deaths worldwide each year. Due to the emergence of drug-resistant strains of *Mycobacterium tuberculosis* (Mtb) there have been significant efforts aimed at developing novel drugs to treat TB. One promising drug target in Mtb is the arabinogalactan biosynthetic enzyme DprE1, and there have been over a dozen unique chemical scaffolds identified which inhibit the activity of this protein. Among the most promising lead compounds are the benzothiazinones BTZ043 and PBTZ169, both of which are currently in or have completed phase IIa clinical trials. Due to the potential clinical utility of these drugs, we sought to identify potential synergistic interactions and new mechanisms of resistance using a genome-scale CRISPRi chemical-genetic screen with PBTZ169. We found that knockdown of *rv0678*, the negative regulator of the *mmpS5/L5* drug efflux pump, confers resistance to PBTZ169. Mutations in *rv0678* are the most common form of resistance to bedaquiline and there is already abundant evidence of these mutations emerging in bedaquiline-treated patients. We confirmed that *rv0678* mutations from clinical isolates confer low level cross-resistance to BTZ043 and PBTZ169. While it is yet unclear whether *rv0678* mutations would render benzothiazinones ineffective in treating TB, these results highlight the importance of monitoring for clinically-prevalent *rv0678* mutations during ongoing BTZ043 and PBTZ169 clinical trials.

## INTRODUCTION

Tuberculosis (TB) is among the most difficult bacterial infections to treat, requiring multiple months of combination therapy to produce a durable cure (1, 2). *Mycobacterium tuberculosis* (Mtb), the causative agent of TB, is intrinsically resistant to many different classes of antimicrobial compounds (3–6). Further, Mtb can evolve acquired drug resistance mutations to every antitubercular drug in clinical use, adding another layer of difficulty to TB treatment (7–9). Current estimates suggest that roughly 5% of global TB cases are caused by multidrug-resistant (MDR) Mtb strains, accounting for roughly 500,000 newly diagnosed, MDR infections each year (2, 10). In certain geographic regions, drug resistance rates can be as high as 40%, highlighting the need for new antitubercular drugs and drug combinations (2).

Drug discovery efforts over the past two decades have identified several Mtb proteins that are commonly found to be the target of new antitubercular compounds (11). Among these “promiscuous targets” are the trehalose dimycolate transporter MmpL3, the cytochrome *bc1* oxidase subunit QcrB, and the decaprenylphosphoryl-β-D-ribofuranose-2′-epimerase subunit DprE1, which is involved in the synthesis of arabinogalactan (11–14). Many distinct chemical scaffolds have been found to inhibit each of these proteins and several of these compounds have advanced to clinical trials (15–19). Among these are the benzothiazinone (BTZ) DprE1 inhibitors including PBTZ169 (20) and BTZ043 (21, 22) which are among the most potent antitubercular compounds discovered thus far, with minimum inhibitory concentrations (MICs) in the low nanomolar range (22). In addition to having potent antitubercular activity, the lead BTZ, PBTZ169, displays synergistic activity with other antitubercular drugs such as bedaquiline, presumably by increasing Mtb cell permeability and bedaquiline uptake (23, 24). PBTZ169 (Macozinone) underwent a phase IIa early bactericidal activity (EBA) clinical trial in Russia where it showed good safety, tolerability and EBA (16, 25). BTZ043 is part of an ongoing phase IIa study in South Africa (18, 26).

Given the clinical promise of this class of drugs, we sought to better understand the bacterial genetic determinants of sensitivity and resistance to PBTZ169. By identifying genes whose inhibition sensitizes Mtb to PBTZ169, we hoped to identify new synergistic drug targets for BTZs. Further, by identifying genes whose inhibition results in PBTZ169 resistance, we hoped to identify novel sources of genetic resistance to BTZs, distinct from the mutations in the drug target which have been well characterized (21, 22, 27).

## RESULTS

### CRISPR interference-based chemical-genetic profiling identifies determinants of PBTZ169 sensitivity and resistance

To identify the genetic determinants of PBTZ169 potency, we performed a genome-scale, chemical-genetic screen using the Mtb CRISPR interference (CRISPRi) platform described previously (28–31). This library contains 96,700 unique single guide RNAs (sgRNAs) targeting over 98% of the genes in the H37Rv Mtb genome. This approach allows for the assessment of chemical-genetic interactions for both *in vitro* essential and non-essential genes (30). The H37Rv CRISPRi library was treated with anhydrotetracycline (ATc) for five days to deplete target proteins, after which the library was split into triplicate cultures treated with either DMSO (vehicle control) or various concentrations of PBTZ169 (**Fig. 1A**). The 0.8 and 1.6 nM PBTZ169 concentrations (**Fig. 1B**) were used for downstream analysis, since these concentrations of PBTZ169 exerted selective pressure without bottlenecking (i.e. sharply reducing complexity) the CRISPRi library. The fitness of each strain was assessed by deep sequencing of the sgRNA encoding region and subsequent gene level analysis was performed using MAGeCK (32).

**Figure 1.**
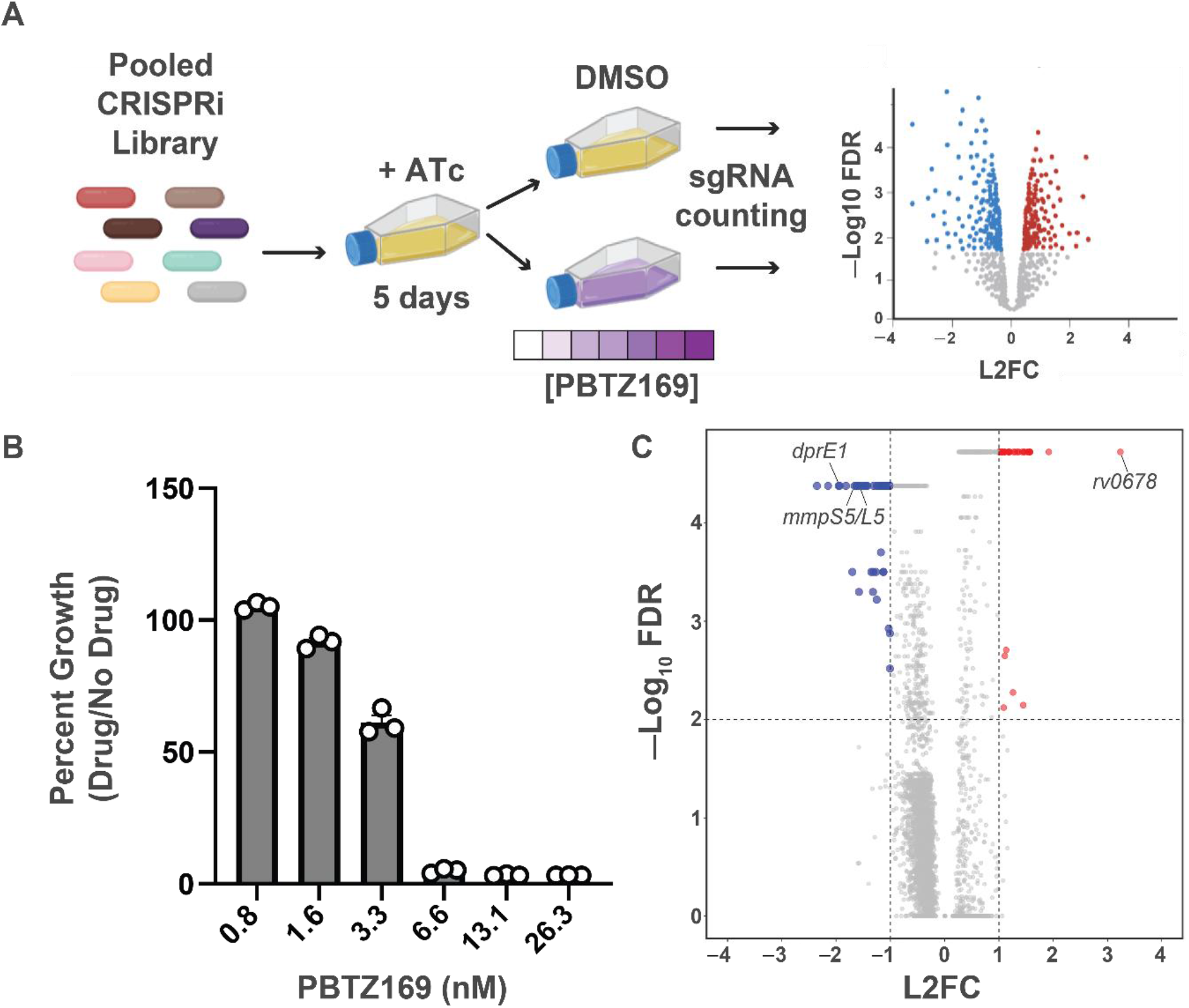
CRISPRi chemical-genetic profiling of PBTZ169. (A) Chemical-genetic profiling workflow. The CRISPRi library contains 96,700 sgRNAs targeting 4,052/4,125 genes in the H37Rv genome. The library was pre-depleted for five days in the presence of anhydrotetracycline (ATc) prior to treatment with DMSO or varying concentrations of PBTZ169 spanning the MIC. After 14 days of library outgrowth, genomic DNA was isolated from all cultures with detectable outgrowth. The sgRNA-encoding region was subjected to deep sequencing for control cultures as well as the 0.8 nM and 1.6 nM PBTZ169 cultures. Hit genes were called by MAGeCK (32). (B) Relative growth of PBTZ169-treated cultures. After 14 days of outgrowth, growth was calculated relative to the DMSO control. Individual replicates are shown. (C) Volcano plot summarizing the log_2_ fold-change (L2FC) value and –Log_10_ of the false discovery rate (FDR) for each gene in the PBTZ169 screen (1.6 nM). Key hit genes are highlighted.

This screen identified a total of 79 chemical-genetic hits (|L2FC>1|, FDR < 0.01; **Fig. 1C, Supplemental file 1**). There were a total of 46 genes for which knockdown resulted in increased PBTZ169 sensitivity and 33 genes for which knockdown resulted in increased PBTZ169 resistance (**Fig. 1C**). Among the most sensitizing hit genes was *dprE1*, consistent with previous studies showing that the genetic knockdown of a drug target generally results in sensitivity to that particular drug (30, 33, 34). Also among the sensitizing hits were a number of thioredoxin genes (*trxC, trxB2, sirA*). Alteration of cellular redox homeostasis could alter

FAD/FADH_2_ levels and make DprE1 more susceptible to PBTZ169-mediated inhibition (21). Further, the screen identified a number of cell wall biosynthetic enzymes (*pbpA, idsA2, mmaA4*) as sensitizing hits. Sensitivity of these knockdown strains to PBTZ169 may reflect the combined effects of inhibiting multiple cell envelope synthetic pathways. Alternatively, these mutants may have a compromised cell envelope, allowing PBTZ169 to more easily reach DprE1 in the mycobacterial periplasm (35, 36). However, the chemical-genetic signature of PBTZ169 is distinct from drugs like rifampicin and vancomycin, for which the envelope appears to be a relevant permeability barrier, suggesting that the activity of PBTZ169 is not limited by drug uptake (30, 37). Finally, we identified the *mmpS5/L5* efflux pump genes as among the most sensitizing hits in the screen. MmpLS5/L5 is a multidrug efflux pump that extrudes a number of antitubercular drugs such as bedaquiline and clofazimine (38). These results suggest that MmpS5/L5 may also be active against PBTZ169.

This chemical-genetic screen was also used to identify BTZ resistance mechanisms independent of drug binding site mutations in DprE1 (22, 27, 39). The strongest resistance hit gene was *rv0678*, which showed a greater than 8-fold enrichment under PBTZ169 treatment conditions compared to the vehicle control (**Fig. 1C, Supplemental file 1**). Rv0678 encodes a transcription factor which negatively regulates the expression of the *mmpS5/L5* efflux pump (38). Loss-of-function mutations in *rv0678* are a common mechanism of resistance to the new antitubercular drug bedaquiline as well as the antileprotic/antitubercular drug clofazimine (38, 40, 41). There are several reports of *rv0678* mutations arising at a high frequency in Mtb clinical isolates (41–45).

Further, some *rv0678* mutations pre-date the introduction of bedaquiline to TB treatment regimens and may have arisen in response to other selective pressures, such as clofazimine treatment of leprosy infections (43). Given the clinical prevalence of *rv0678* mutations as well as the strong *rv0678* signature in the PBTZ169 screen, we sought to determine whether this may be a relevant mechanism of resistance to BTZs.

### Clinically prevalent mutations in *rv0678* confer low-level resistance to BTZs

In order to assess whether mutations in *rv0678* may confer cross-resistance to BTZs, we first isolated a select group of three *rv0678* mutants associated with clinical bedaquiline resistance (42–44, 46) using single-stranded DNA recombineering in H37Rv Mtb. Mutations in *rv0678* were confirmed by Sanger sequencing and strains were subjected to whole genome sequencing to ensure that there were no other mutations that are likely to influence drug susceptibility. We tested the susceptibility of all three *rv0678* mutants and the wild-type strain to sixteen different antitubercular drugs (**Fig. 2, Fig. S1, Table 1**). Consistent with the CRISPRi screening results, the *rv0678* frameshift mutants showed a ∼4-fold increase in IC_50_ to PBTZ169 and a ∼3-fold increase in IC_50_ to BTZ043. The C268T missense mutant showed slightly smaller IC_50_ shifts of 2.4-fold and 1.7-fold to PBTZ169 and BTZ043, respectively. These IC_50_ shifts were slightly smaller than what was seen for bedaquiline but comparable to what was seen for clofazimine. The *rv0678* mutants showed nearly identical susceptibility to the other drugs tested, including two chemically-distinct DprE1 inhibitors, TCA-1 and IN-2 (47, 48). The lack of cross-resistance to TCA-1 and IN-2 suggest that MmpL5/S5 efflux is not relevant for all DprE1 inhibitors. Interestingly, *rv0678* mutants also showed resistance to fusidic acid, an antibiotic not used in TB treatment but used widely to treat skin and soft tissue infections caused by *Staphylococcus aureus* (49). These results suggest that orally-administered fusidic acid may be a potential selective pressure for Mtb *rv0678* mutations in areas where bedaquiline and clofazimine have not been introduced (43).

**Figure 2.**
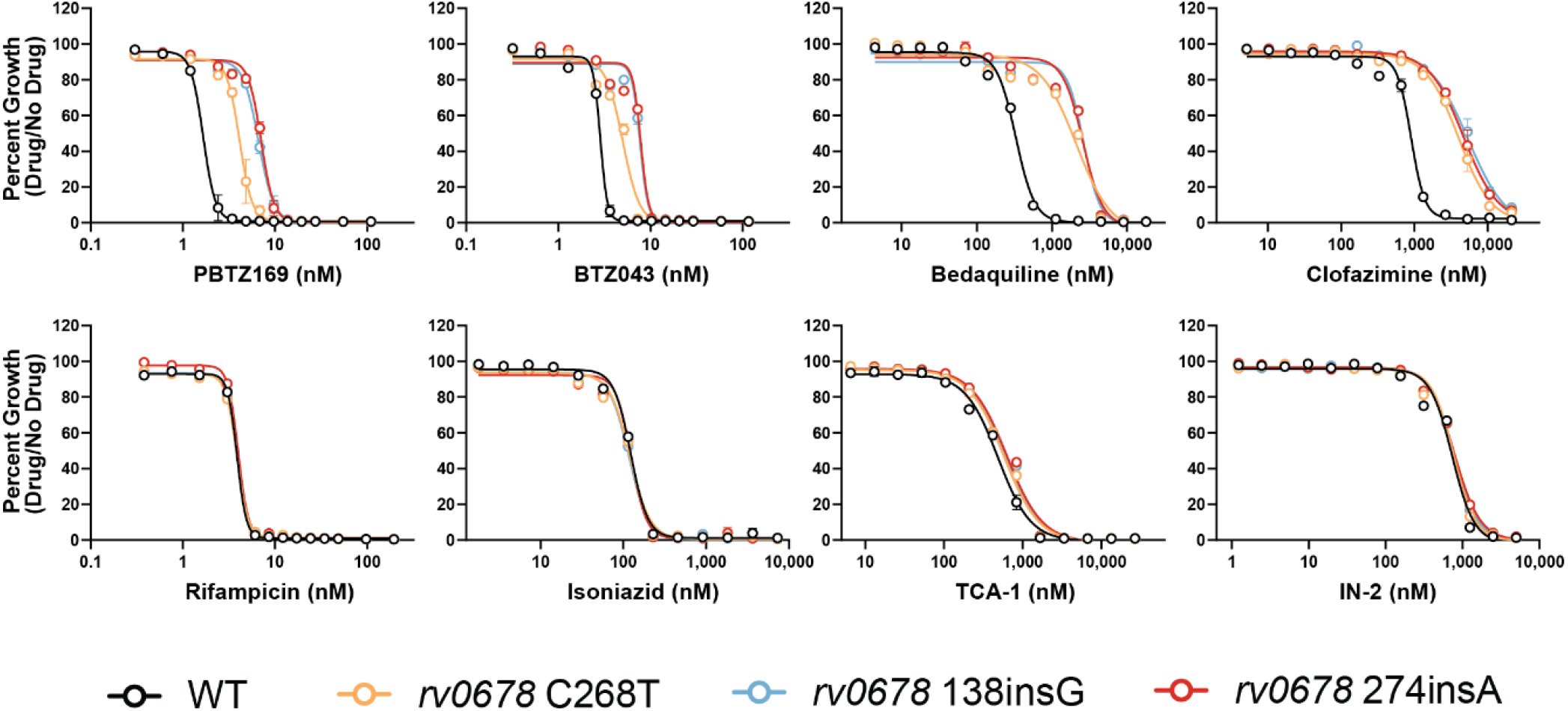
Antibiotic susceptibility of *rv0678* mutants. Dose-response curves (mean ± s.e.m., n= 3 replicates) for the indicated Mtb strains. Results are representative of two independent experiments.

**Table 1:** ^a^Susceptibility (IC_50_) of *rv0678* mutants to 16 antibiotics

| Antibiotic | Target process | Wild-type | *rv0678* 138insG | *rv0678* 274insA | *rv0678* C268T | IC<sub>50</sub> ratio<sup>b</sup> |
| --- | --- | --- | --- | --- | --- | --- |
| PBTZ169 | Arabinogalactan | 1.7 | 6.6 | 7.1 | 4.1 | 4.2 |
| BTZ043 | Arabinogalactan | 2.9 | 7.7 | 7.8 | 5 | 2.7 |
| Bedaquiline | ATP synthase | 330 | 2700 | 2600 | 2100 | 7.9 |
| Clofazimine | Respiration | 900 | 5400 | 4700 | 4000 | 5.2 |
| Rifampicin | Transcription | 3.9 | 3.9 | 4 | 3.9 | 1.0 |
| Isoniazid | Mycolic acid | 120 | 120 | 120 | 120 | 1.0 |
| TCA-1 | Arabinogalactan | 480 | 610 | 630 | 570 | 1.3 |
| IN-2 | Arabinogalactan | 730 | 770 | 780 | 760 | 1.1 |
| Ethambutol | Arabinogalactan | 2400 | 2800 | 3200 | 3100 | 1.3 |
| Levofloxacin | DNA replication | 700 | 700 | 700 | 690 | 1.0 |
| Pretomanid | Mycolic acid | 360 | 400 | 430 | 440 | 1.2 |
| Linezolid | Translation | 1300 | 1400 | 1300 | 1200 | 1.0 |
| Streptomycin | Translation | 420 | 410 | 430 | 440 | 1.0 |
| Amikacin | Translation | 930 | 900 | 950 | 980 | 1.0 |
| Capreomycin | Translation | 1200 | 1200 | 1200 | 1200 | 1.0 |
| Fusidic Acid | Translation | 4200 | 26000 | 26000 | 17000 | 6.2 |

<sup>a</sup> IC<sub>50</sub> values reflect drug concentrations in nanomolar (nM)
<sup>b</sup> The IC<sub>50</sub> ratio was calculated by dividing the IC<sub>50</sub> for the *rv0678* 274insA mutant by the IC<sub>50</sub> for the wild-type strain.

Given the phenotypes we observed with *rv0678* mutants, we next sought to confirm that overexpression of the MmpS5/L5 efflux pump is responsible for the BTZ resistance in *rv0678* mutants. To do this, we made use of a lineage 1 clinical isolate of Mtb harboring a nonsense mutation (Tyr300Stop) in *mmpL5* (50). This strain was complemented with a functional copy of the *mmpS5/L5* operon expressed from the native promoter to restore wild-type expression or by the *hsp60* promoter to overexpress the efflux pump (30). Consistent with the previous results, overexpression of *mmpS5/L5* resulted in increased IC_50_ for PBTZ169, BTZ043, bedaquiline, clofazimine, and fusidic acid, but not the other drugs tested (**Fig. 3, Fig. S2, Table 2**). Interestingly, while the parental *mmpL5* Tyr300Stop mutant was hypersusceptible to bedaquiline, this strain was only modestly more sensitive to the BTZs, clofazimine, and fusidic acid, consistent with previous studies where gain-of-function and loss-of-function mutations may not necessarily produce opposing phenotypes of equal magnitude (51, 52).

**Figure 3.**
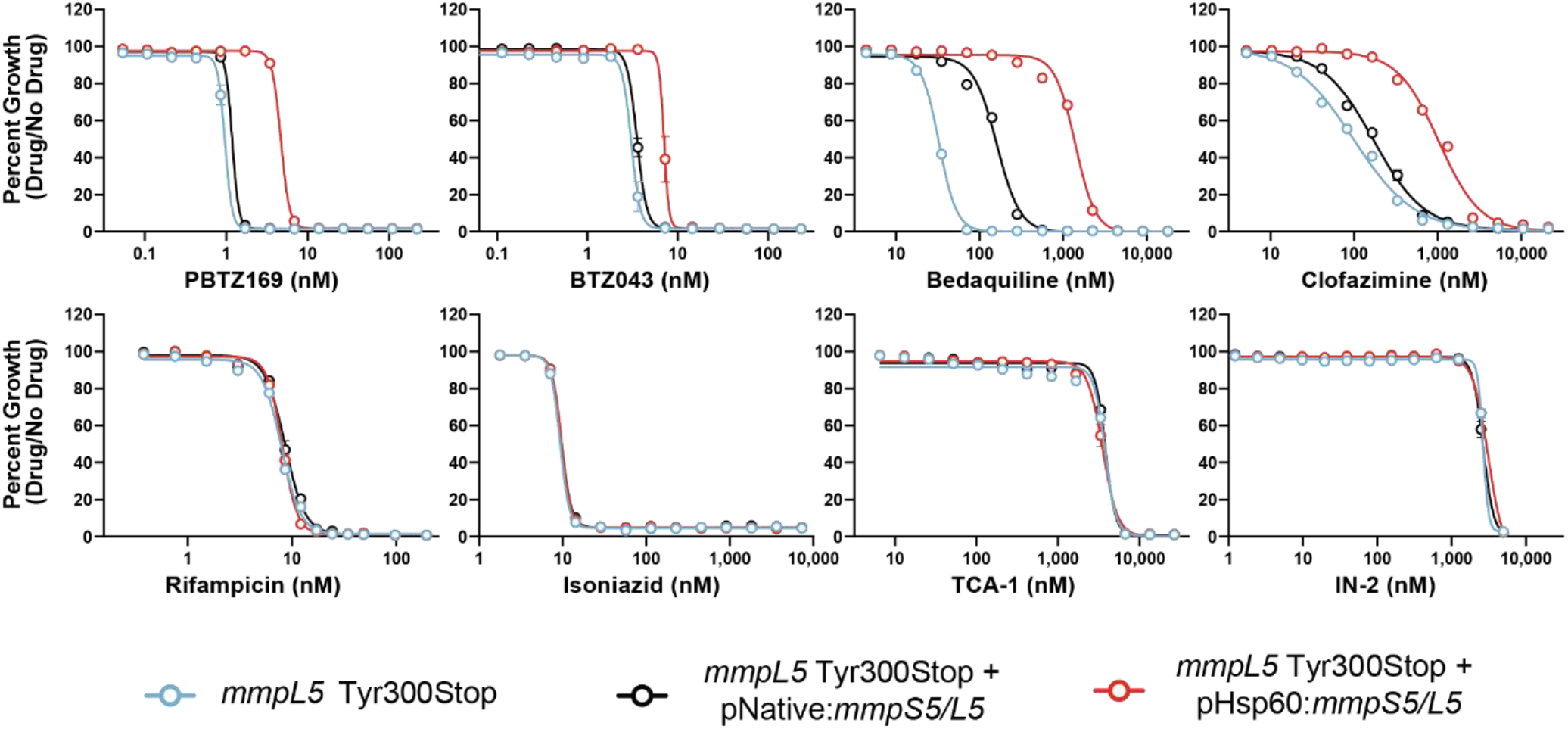
Antibiotic susceptibility of *mmpL5* loss-of-function and gain-of-function strains. Dose-response curves (mean ± s.e.m., n= 3 replicates) for the indicated Mtb strains. Results are representative of two independent experiments. The Tyr300Stop mutant (blue) harbors an empty version of the *mmpS5/L5* complementation vector (30).

**Table 2:** ^a^Susceptibility of *mmpL5* loss-of-function and gain-of-function strains to 16 antibiotics

| Antibiotic | Target process | *mmpL5* Tyr300Stop | *mmpL5* Tyr300Stop + pNative:*mmpS5/L5* | *mmpL5* Tyr300Stop + pHsp60:*mmpS5/L5* | IC<sub>50</sub> ratio<sup>a</sup> |
| --- | --- | --- | --- | --- | --- |
| PBTZ169 | Arabinogalactan | 1.0 | 1.2 | 4.7 | 3.9 |
| BTZ043 | Arabinogalactan | 3.1 | 3.5 | 7 | 2 |
| Bedaquiline | ATP synthase | 33 | 160 | 1400 | 8.8 |
| Clofazimine | Respiration | 100 | 170 | 1030 | 6.1 |
| Rifampicin | Transcription | 7.2 | 7.8 | 7.3 | 0.9 |
| Isoniazid | Mycolic acid | 9.3 | 9.7 | 9.7 | 1.0 |
| TCA-1 | Arabinogalactan | 3800 | 3800 | 3500 | 0.9 |
| IN-2 | Arabinogalactan | 2700 | 2700 | 3100 | 1.1 |
| Ethambutol | Arabinogalactan | 1900 | 1900 | 2300 | 1.2 |
| Levofloxacin | DNA replication | 900 | 960 | 970 | 1.0 |
| Pretomanid | Mycolic acid | 750 | 810 | 830 | 1.0 |
| Linezolid | Translation | 1100 | 1100 | 1200 | 1.1 |
| Fusidic Acid | Translation | 1800 | 3400 | 15000 | 4.4 |

<sup>a</sup> IC<sub>50</sub> values reflect drug concentrations in nanomolar (nM)
<sup>b</sup> The IC<sub>50</sub> ratio was calculated by dividing the IC<sub>50</sub> for the pHsp60-driven *mmpS5/L5* strain by the IC<sub>50</sub> for the pNative-driven *mmpS5/L5* strain.

## DISCUSSION

BTZ DprE1 inhibitors are promising antitubercular agents that are currently in early stage clinical trials (16, 17, 53). A better understanding of the genetic factors that influence the potency of these drugs could help to inform the design of synergistic drug combinations and to diagnose the emergence of BTZ resistance. Here we applied a genome-scale CRISPRi screen to identify the determinants of sensitivity and resistance to PBTZ169. We identified dozens of hit genes, a number of which are essential, which may be targeted to potentiate the activity of BTZs. Target-based drug discovery efforts towards these proteins may identify synergistic partners for use with BTZs. Future chemical-genetic efforts could apply a similar screening pipeline to study other non-BTZ DprE1 inhibitors (53, 54) to identify common and unique components of their chemical-genetic signatures. This may provide a global overview of the genes and pathways which restrict the activity of these compounds and allow for the design of optimized DprE1 inhibitors that fully exploit this chemically vulnerable drug target.

In addition to finding strategies to potentiate the activity of PBTZ169, we hoped to use this screen to identify resistance mechanisms for BTZs that are mediated by loss-of-function mutations. Interestingly, the strongest enriched hit gene found in our screen was *rv0678*, which was chosen for follow-up investigation due to the potential clinical relevance of this finding. We observed that common *rv0678* mutations from Mtb clinical isolates confer low-level resistance to PBTZ169 and BTZ043. Resistance is also observed in Mtb strains directly overexpressing the *mmpS5/L5* operon, suggesting that efflux of BTZs is the relevant mechanism of resistance. Consistent with our findings, another recent study isolated PBTZ169-resistant mutants on progressively higher drug concentrations and identified a number of clones harboring mutations in *rv0678* (55). Although the authors do not test the individual impact of each *rv0678* mutation, the results presented here suggest that these are likely to be *bona fide*, low-level PBTZ169 resistance mutations.

Importantly, *rv0678* mutations did not promote resistance to the chemically distinct (47, 48) DprE1 inhibitors TCA-1 and IN-2. This observation argues against a role for perturbed Mtb physiology as a result of elevated MmpL5/S5 activity in promoting resistance to BTZs, rather suggesting efflux is the relevant resistance mechanism. Previous work has convincingly argued that DprE1 localizes to the periplasm, suggesting that periplasmic localization contributes to the chemical vulnerability of DprE1: inhibitors need not cross the Mtb plasma membrane to the cytosol in order to engage their target (35, 36). If this is true, how might the MmpL5/S5 efflux pump, which is localized to the plasma membrane, promote resistance to BTZs? There are several potential explanations to these findings. First, it is possible that BTZs can covalently inhibit DprE1 in the cytosol prior to Tat-mediated transport of the folded DprE1 to the periplasm. While experimental evidence for Tat-mediated DprE1 export is lacking, DprE1 is predicted to encode a Tat signal peptide (56, 57). In this scenario, MmpL5/S5 efflux of BTZs from the cytosol will hinder cytoplasmic DprE1 inhibition. Second, it is possible that MmpL5/S5 could efflux BTZs out of the periplasm to prevent DprE1 engagement, through a mechanism similar to that of the AcrB efflux pump in *E. coli* (58). This would presumably require MmpL5/S5 interaction with mycomembrane spanning proteins to efflux BTZs outside of the Mtb envelope (59–61). Along these lines, studies in Gram-negative bacteria have shown that efflux pumps can form large protein complexes that span both bacterial membranes and promote resistance to beta-lactam antibiotics, whose PBP targets also localize to the periplasm (58, 59, 62–66).

Although the exact mechanism of MmpS5/L5-mediated BTZ resistance is not clear, these results suggest that *rv0678* mutations could limit the clinical efficacy of BTZs, especially in areas where bedaquiline has already been used. While the BTZ series are remarkably potent, murine pharmacokinetic data suggests that PBTZ169 accumulates to concentrations close to its MIC, particularly in caseous lesions (53). The mouse data suggest that small increases in MIC (67) may be sufficient to render PBTZ169 less effective. Future studies are necessary to test whether *rv0678* mutations could contribute to BTZ treatment failure in humans.

## ACKNOWLEDGMENTS

We thank members of the Rock laboratory, Véronique Dartois, Katarína Mikušová, Valerie Mizrahi, Max O’Donnell, Michelle Larsen, and James Millard for comments on the manuscript and/or helpful discussions as well as Xing Wang, Jenny Xiang, and Adrian Tan of the Weill Cornell Genomics Core for NGS. This work was supported by the Potts Memorial Foundation (M.A.D.), the Department of Defense (PR192421, J.M.R.), the Robertson Therapeutic Development Fund (J.M.R.), and an NIH/NIAID New Innovator Award (1DP2AI144850-01, J.M.R.). The illustration in figure 1A was generated using BioRender software.

## AUTHOR CONTRIBUTIONS

Conceptualization: N.C.P. and J.M.R. Investigation: N.C.P. Data Analysis: N.C.P., Z.A.A., and M.A.D. Writing—original draft: N.C.P. and J.M.R. Writing—review and editing: N.C.P., Z.A.A., M.A.D., and J.M.R. Funding acquisition: J.M.R. Supervision: J.M.R.

## SUPPLEMENTAL FIGURES

**Supplemental Figure 1.**
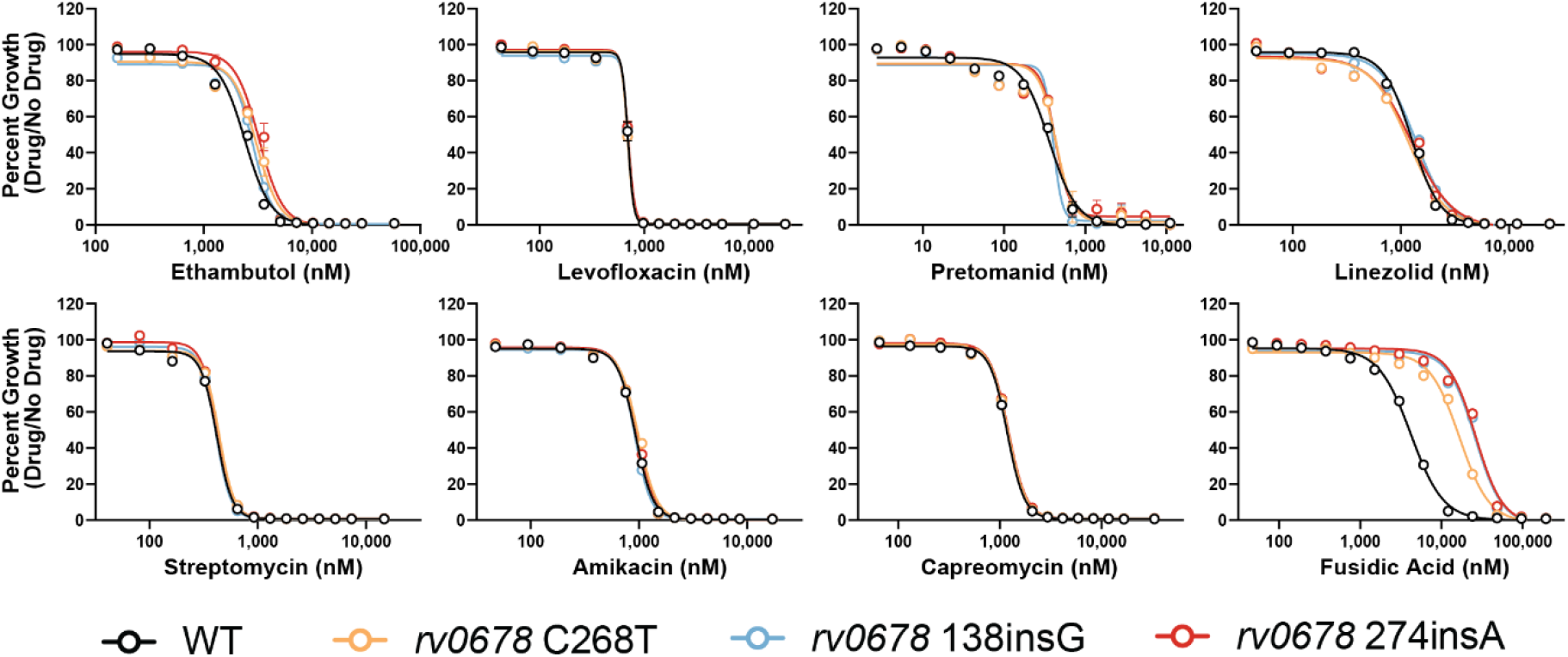
Antibiotic susceptibility of *rv0678* mutants. Dose-response curves (mean ± s.e.m., n= 3 replicates) for the indicated Mtb strains. Results are representative of two independent experiments.

**Supplemental Figure 2.**
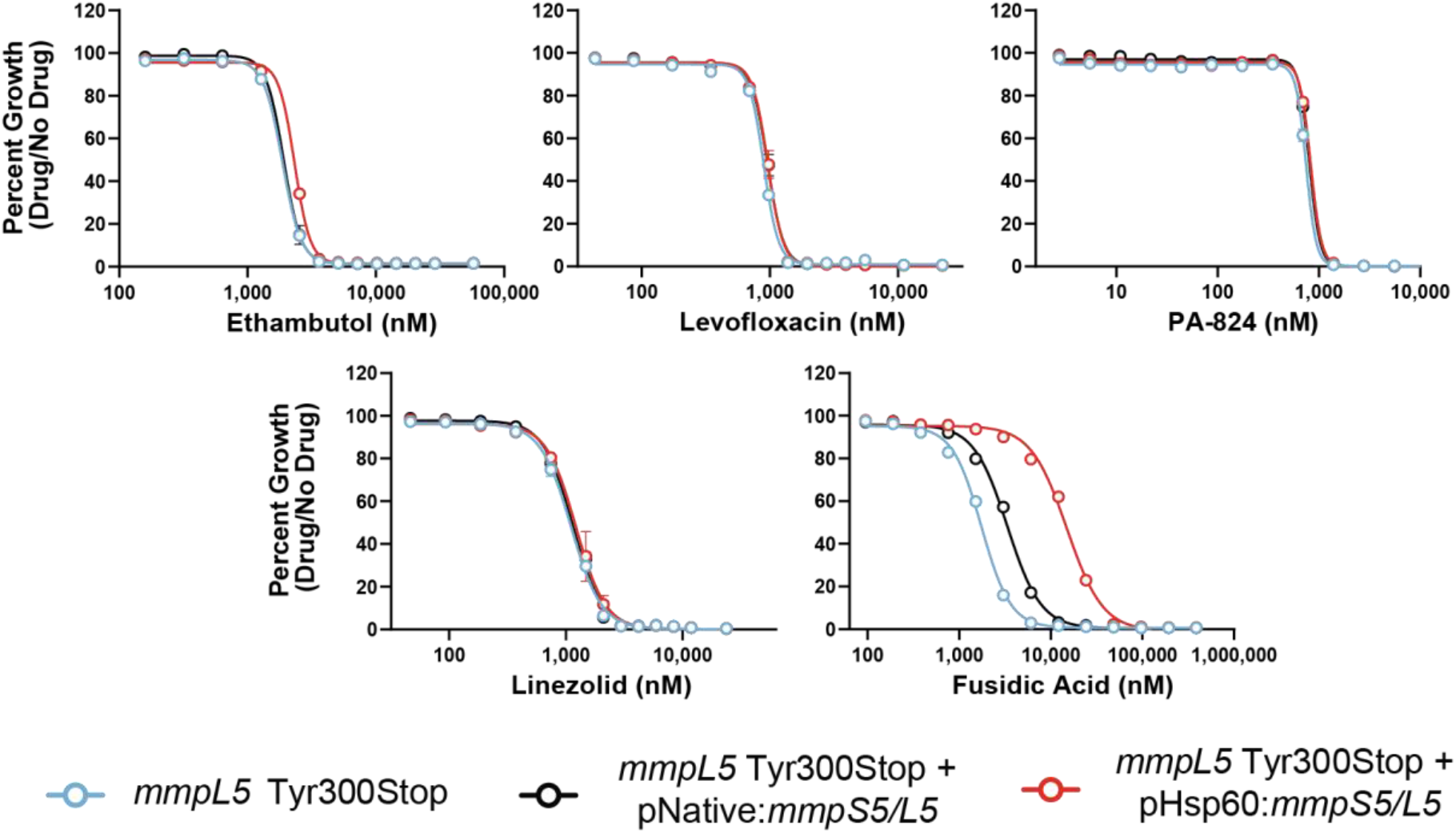
Antibiotic susceptibility of *mmpL5* loss-of-function and gain-of-function strains. Dose-response curves (mean ± s.e.m., n= 3 replicates) for the indicated Mtb strains. Results are representative of two independent experiments.

## MATERIALS AND METHODS

### Mycobacterial cultures

Mtb was grown at 37°C in Difco Middlebrook 7H9 broth or on 7H10 agar supplemented with 0.2% glycerol (7H9) or 0.5% glycerol (7H10), 0.05% Tween-80, 1X oleic acid-albumin-dextrose-catalase (OADC) and the appropriate antibiotics (kanamycin 10-20 μg/mL or zeocin 20μg/mL). Anhydrotetracycline (ATc) was used at 100ngml−1 in the CRISPRi screen and at 500 ng/mL for ssDNA recombineering (68). Mtb cultures were grown standing in tissue culture flasks (unless otherwise indicated) with 5% CO_2_. Mtb strains are derivatives of the H37Rv background, except those shown in Fig. 3 and supplemental Fig. 2. Those strains are derivatives of a Lineage 1.1 strain (ITM-2018-00084) from the Belgian Coordinated Culture Collection (BCCM) (50). This strain harbors the Tyr300Stop mutation in *mmpL5*. The *mmpS5/L5* complementation and overexpression strains are described by Li et al. (30).

### Pooled CRISPRi chemical-genetic screening

Chemical-genetic screens were initiated by thawing 2 × 1ml aliquots (1.0 OD_600_ units/mL) of the Mtb CRISPRi library (RLC12; Addgene 163954) and inoculating each aliquot into 19ml 7H9 supplemented with kanamycin (10 μg/mL) in a vented tissue culture flask (T-75; Falcon 353136)(30). The starting OD_600_ of each culture was approximately 0.05. Cultures were expanded to OD_600_=1.0, pooled and passed through a 10 μm cell strainer (pluriSelect 43-50010-03) to obtain a single-cell suspension. The single-cell suspension was then back-diluted to an OD_600_ of 0.05 in 4 × 25 mL cultures and treated with ATc (100 ng/ml final concentration) to initiate target pre-depletion. After 5 days of pre-depletion triplicate cultures were then inoculated at OD_600_ of 0.05 in 10ml 7H9 supplemented with ATc (100ng/mL), kanamycin (10 μg/mL), and the indicated PBTZ169 concentration or a DMSO vehicle control. In all cultures the final DMSO concentration was normalized to 0.6%. Pooled CRISPRi chemical-genetic screens were performed in vented tissue culture flasks (T-25; Falcon 353109) under standing (non-shaking) conditions. Cultures were outgrown for 14d at 37 °C, 5% CO2. ATc was replenished at 100ngml−1 at day 7. After 14d outgrowth, OD_600_ values were measured for all cultures to empirically determine the MIC for each drug. The lowest three concentrations of PBTZ169-treatment were subjected to sequencing. However, the 3.3 nM condition showed evidence of library bottlenecking following sgRNA sequencing, likely because this condition was near the MIC. Downstream analysis was performed on the 0.8 nM and 1.6 nM conditions.

### Genomic DNA extraction and library preparation for Illumina sequencing

Genomic DNA was isolated from bacterial pellets using the CTAB−lysozyme method as previously described (28). After drug treatment 10-20 OD_600_ units of the cultures were pelleted by centrifugation (10 minutes at 4,000xg) and were resuspended in 1ml PBS +0.05% Tween-80. Cell suspensions were centrifuged again for 5min at 4,000×g, the supernatant was removed, and pellets were frozen until processing. Upon thawing, pellets were resuspended in 800 µl TE buffer (10mM Tris pH 8.0, 1mM EDTA) + 15mg/mL lysozyme (Alfa Aesar J60701-06) and incubated at 37 °C for 16h. Next, 70µl 10% SDS (Promega V6551) and 5 µl proteinase K (20 mg/mL, Thermo Fisher 25530049) were added and samples were incubated at 65°C for 30min. Subsequently, 100 µl 5M NaCl and 80 µl 10% CTAB (Sigma Aldrich H5882) were added, and samples were incubated for an additional 30min at 65 °C. Finally, 750 µl ice-cold chloroform was added and samples were mixed. After centrifugation at 16,100×g and extraction of the aqueous phase, samples were removed from the biosafety level 3 facility. Samples were then treated with 25 µg RNase A (Bio Basic RB0474) for 30min at 37 °C, followed by extraction with phenol:chloroform:isoamyl alcohol (pH 8.0, 25:24:1, Thermo Fisher BP1752I-400), then chloroform. Genomic DNA was precipitated from the final aqueous layer (600 µl) with the addition of 10 µl 3M sodium acetate and 360 µl isopropanol. DNA pellets were spun at 21,300×g for 30min at 4 °C and washed 2× with 750 µl 80% ethanol. Pellets were dried and resuspended with elution buffer (Qiagen 19086) before spectrophotometric quantification. The concentration of isolated genomic DNA was quantified using the DeNovix dsDNA high sensitivity assay (KIT-DSDNA-HIGH-2; DS-11 series spectrophotometer/fluorometer).

Next generation sequencing was performed as described by Bosch et al (28). The sgRNA-encoding region was amplified from 500ng genomic DNA with 17 cycles of PCR using NEBNext Ultra II Q5 master mix (NEB M0544L) as described in. Each PCR reaction contained a pool of forward primers (0.5μM final concentration) and a unique indexed reverse primer (0.5μM). Forward primers contain a P5 flow cell attachment sequence, a standard Read1 Illumina sequencing primer binding site and custom stagger sequences to guarantee base diversity during Illumina sequencing. Reverse primers contain a P7 flow cell attachment sequence, a standard Read2 Illumina sequencing primer binding site and unique barcodes to allow for sample pooling during deep sequencing. Following PCR amplification, each ∼230bp amplicon was purified using AMPure XP beads (Beckman–Coulter A63882) using one-sided selection (1.2×). Bead-purified amplicons were further purified on a Pippin HT 2% agarose gel cassette (target range 180–250bp; Sage Science HTC2010) to remove carry-over primer and genomic DNA. Eluted amplicons were quantified with a Qubit 2.0 fluorometer (Invitrogen), and amplicon size and purity were quality controlled by visualization on an Agilent 2100 bioanalyzer (high sensitivity chip; Agilent Technologies 5067–4626). Next, individual PCR amplicons were multiplexed into 10nM pools and sequenced on an Illumina sequencer according to the manufacturer’s instructions. To increase sequencing diversity, a PhiX spike-in of 2.5–5% was added to the pools (PhiX sequencing control v3; Illumina FC-110-3001). Samples were run on the Illumina NovaSeq 6000 platform (single-read 1 ×85 cycles and 6 × i7 index cycles).

### Isolation of *rv0678* mutants by single-stranded DNA recombineering

The 138insG frameshift allele was observed in a bedaquiline-treated patient and emerged as the dominant clone over the course of treatment (41). Another frameshift mutation, 274insA, was identified multidrug-resistant TB patients with no prior evidence of bedaquiline or clofazimine treatment (43). The C268T missense mutation, which results in the Arg90Cys amino acid change, has been detected in MDR-TB patients from South Africa and Korea (42, 44) Single-stranded DNA recombineering was performed as described by Murphy et al (68). Briefly, H37Rv Mtb was transformed with the episomal RecT recombinase-expressing plasmid pKM402 (Addgene plasmid # 107770) and was selected on 7H10 agar containing kanamycin (20 µg/mL). A single colony was picked and grown up in 7H9 + kanamycin (20 µg/mL). A single frozen stock of the RecT strain was thawed and expanded to 50 mL in 7H9 + kanamycin (20 µg/mL) in Erlenmeyer flasks under shaking conditions (37°C). After reaching an OD_600_ of 0.8, anhydrotetracycline (ATc) was added at a final concentration of 500 ng/mL to induce RecT expression. After 8 more hours of shaking incubation, sterile glycine was added to a final concentration of 200 mM to improve DNA uptake. After an additional 16 hours of shaking incubation, cells were pelleted (4,000xg) and washed a total of three times in sterile 10% glycerol. On the final wash, cells were resuspended in 5 mL of 10% glycerol. For each transformation, 400 µL of electrocompetent cells was mixed with 1 µg each ssDNA oligo of interest (see supplemental table 1). Cells were electroporated using the Gene Pulser X cell electroporation system (Bio-Rad 1652660) set at 2,500V, 1,000Ω and 25μF and were recovered for 24 hours in 15 mL of fresh 7H9 under 37°C, shaking conditions. 200 µL of each recovered culture was plated on bedaquiline-containing 7H10 plates (125 ng/mL). The control strain shown in Fig. 2 and supplemental Fig. 1 was electroporated with no DNA and plated on antibiotic-free 7H10 plates. Plates were incubated for 28 days, after which colonies were picked and grown in 7H9 + bedaquiline at 500 ng/mL, due to the higher MIC of bedaquiline in liquid culture. For each strain of interest, a total of 4 colonies were picked and genomic DNA was extracted as described above. *rv0678* was amplified and sanger sequenced. Successful recombinants were then submitted to whole genome sequencing (WGS) via Migs (Microbial Genome Sequencing Center, Pittsburgh PA). Whole genome sequences were derived from FASTQ files and any mutations were identified using Snippy(69). WGS results confirmed the presence of the desired *rv0678* mutations and absence of any other mutations known to influence drug susceptibility. The negative control “wild-type” strain contained no mutations in *rv0678* or other genes known to influence drug susceptibility.

### Antibacterial activity measurements

All compounds (see supplemental table 2) were dissolved in DMSO (VWR V0231) and dispensed using an HP D300e digital dispenser in a 384-well plate format using a 2-fold dilution series. For certain drugs, concentrations near the MIC reflect a 1.41-fold dilution series to provide higher resolution. DMSO did not exceed 1% of the final culture volume and was maintained at the same concentration across all samples. Cultures were growth synchronized to late logarithmic phase (∼ OD_600_ = 0.8) and then back-diluted to a starting OD_600_ of 0.01. 50 µl cell suspension was plated in triplicate in wells containing the test compound. Plates were incubated standing at 37 °C with 5% CO2. OD_580_ was evaluated using a Tecan Spark plate reader at 10–11 days post-plating and percent growth was calculated relative to the DMSO vehicle control for each strain. IC_50_ measurements were calculated using a nonlinear fit in GraphPad Prism. For all MIC curves, data represent the mean ± s.e.m. for triplicates. Data are representative of two independent experiments.

## SUPPLEMENTAL MATERIALS

**Supplemental table 1:**
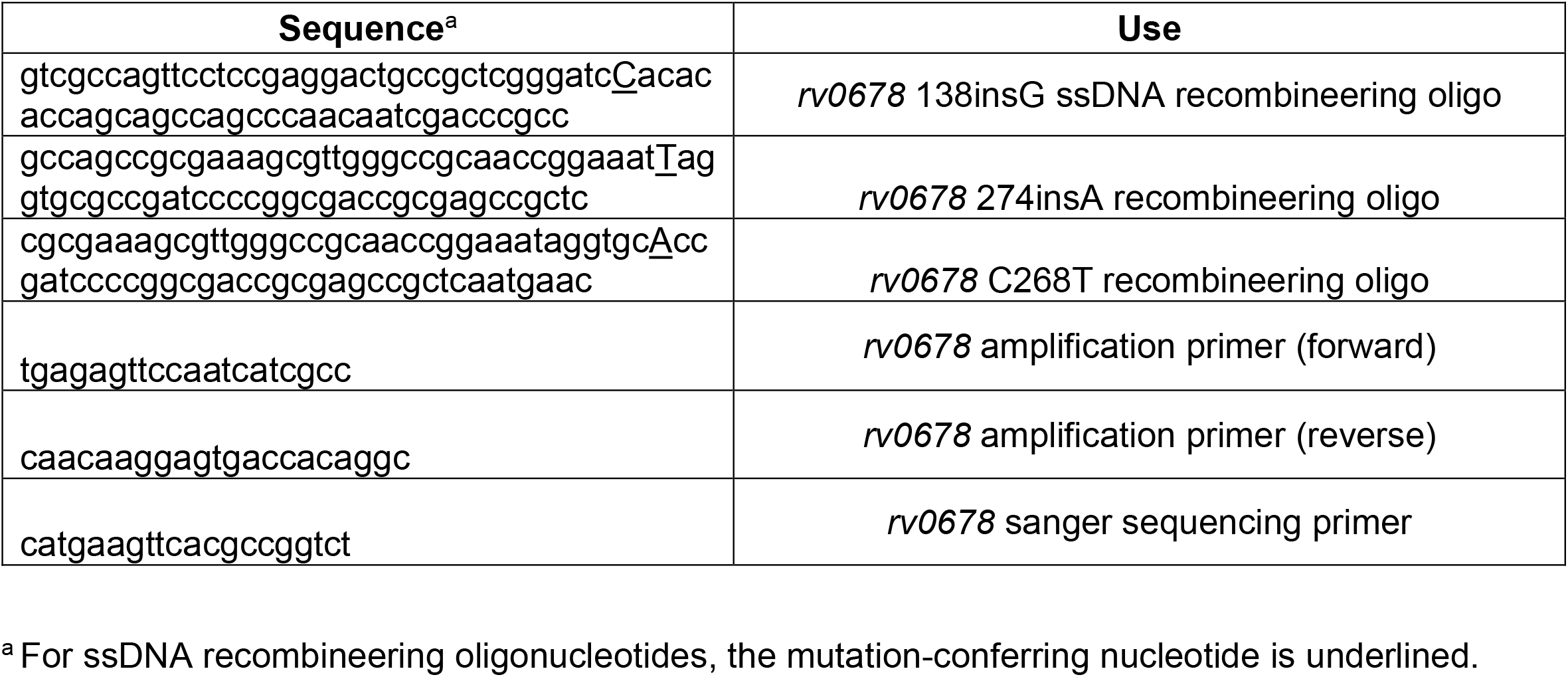
Oligonucleotides used in this study

**Supplemental table 2:** Antimicrobial compounds used in this study

| Antibiotic | Product number |
| --- | --- |
| Amikacin | A0365900 (Sigma Aldrich) |
| Anhydrotetracycline | AC233135000 (Fisher Scientific) |
| Bedaquiline | A12327-5 (AdooQ Biosciences) |
| BTZ043 | 22930 (Cayman Chemical) |
| Capreomycin Sulfate | PHR1716 (Sigma Aldrich) |
| Clofazimine | TCC2866 (VWR) |
| Ethambutol dihydrochloride | E4630 (Sigma Aldrich) |
| Fusidic acid sodium salt | F0881 (Sigma Aldrich) |
| IN-2 DprE1 inhibitor (compound 18(47)) | HY-100531 (MedChem Express) |
| Isoniazid | I3377 (Sigma Aldrich) |
| Kanamycin | K-120-50 (GoldBio) |
| Levofloxacin | 28266 (Sigma Aldrich) |
| Linezolid | SML1290 (Sigma Aldrich) |
| PBTZ169 | 22202 (Cayman Chemical) |
| Rifampicin | R0079 (TCI) |
| Streptomycin sulfate | S-150-100 (GoldBio) |
| TCA-1 (DprE1 inhibitor) | 50-589-30001 (Fisher Scientific) |
| Zeocin | J67140-8EQ (Alfa Aesar) |

